# The Neuron-specific IIS/FOXO Transcriptome in Aged Animals Reveals Regulatory Mechanisms of Cognitive Aging

**DOI:** 10.1101/2023.07.28.550894

**Authors:** Yifei Weng, Shiyi Zhou, Katherine Morillo, Rachel Kaletsky, Sarah Lin, Coleen T. Murphy

## Abstract

Cognitive decline is a significant health concern in our aging society. Here, we used the model organism *C. elegans* to investigate the impact of the IIS/FOXO pathway on age-related cognitive decline. The *daf-2* Insulin/IGF-1 receptor mutant exhibits a significant extension of learning and memory span with age compared to wild-type worms, an effect that is dependent on the DAF-16 transcription factor. To identify possible mechanisms by which aging *daf-2* mutants maintain learning and memory with age while wild-type worms lose neuronal function, we carried out neuron-specific transcriptomic analysis in aged animals. We observed downregulation of neuronal genes and upregulation of transcriptional regulation genes in aging wild-type neurons. By contrast, IIS/FOXO pathway mutants exhibit distinct neuronal transcriptomic alterations in response to cognitive aging, including upregulation of stress response genes and downregulation of specific insulin signaling genes. We tested the roles of significantly transcriptionally-changed genes in regulating cognitive functions, identifying novel regulators of learning and memory. In addition to other mechanistic insights, comparison of the aged vs young *daf-2* neuronal transcriptome revealed that a new set of potentially neuroprotective genes is upregulated; instead of simply mimicking a young state, *daf-2* may enhance neuronal resilience to accumulation of harm and take a more active approach to combat aging. These findings suggest a potential mechanism for regulating cognitive function with age and offer insights into novel therapeutic targets for age-related cognitive decline.

## Introduction

The loss of cognitive function is a rising problem in our aging society. A 2008 study estimated that at least 22.2% (about 5.4 million) individuals over the age of 71 in the United States have at least mild cognitive impairment^1–3^. Furthermore, global dementia cases are predicted to triple from an estimated 57.4 million cases in 2019 to 152.8 million cases in 2050^4,5^. As most industrialized countries are experiencing a rapid increase in the proportion of the aged population, understanding and potentially preventing the underlying issues of neuronal structural and behavioral decline associated with aging is crucial for societal health.

*C. elegans* is an excellent model system for studying neuronal aging, given its tractable genetics, short lifespan, and simple nervous system^6^. Most importantly, *C. elegans* experiences rapid loss of learning and memory with age^7^: by Day 4 of adulthood, all long-term associative memory ability is lost, and by Day 8, *C. elegans* cannot carry out associative learning or short-term associative memory^7^ – despite the fact that these worms can still move and chemotaxis perfectly well. That is, with age worms first lose long-term memory ability (by Day 4), then short-term memory and learning ability (Day 6-8), then chemotaxis (Day 10-12), then motility (Day 16)^7,8^. Because learning and memory decline extremely early, we consider worms that are only a week old to already be “cognitively aged,” despite the fact that they can chemotax and move well, and will continue to live for another one to two weeks. Therefore, we can examine neurons from these 7-8 day-old adults to explore the causes of these cognitive declines in animals that are otherwise quite healthy. Many human neuronal aging phenotypes and genes of interest for mammalian neuronal function are conserved in *C. elegans*^9^, making discoveries in *C. elegans* possibly applicable to humans.

The Insulin/IGF-1-like signaling (IIS)/FOXO pathway was first discovered to play a role in longevity in *C. elegans*. The lifespan of *daf-2*/Insulin/IGF-1 receptor mutants is twice that of wild-type animals^10^, and this lifespan extension requires the downstream Forkhead box O (FOXO) transcription factor DAF-16^10^. DAF-16/FOXO controls the expression of many genes that contribute to longevity, including stress response, proteostasis, autophagy, antimicrobial, and metabolic genes^11^. As a conserved regulator, the IIS/FOXO pathway also regulates longevity in Drosophila, mice, and humans^12–15^. In addition to regulating lifespan, the IIS pathway regulates neuronal function via the FOXO transcription factor. In particular, *C. elegans* IIS/*daf-2* mutants display DAF-16-dependent improved learning, short-term memory, and long-term memory^7^. While both young and old *daf-2* adult worms display increased learning and memory relative to wild-type, the duration of this extension is not known, and the mechanisms by which *daf-2* mutants maintain neuronal function in older worms is not yet understood. Compared to wild-type worms, *daf-2* mutants better maintain maximum velocity^8^, motility^16,17^, neuromuscular junctions, the ability to regenerate axons^18,19^, and neuronal morphology with age^20–22^. In particular, we previously showed that while *daf-2* has lower observed motility on food^23^, this apparent is due to its high levels of a food receptor, ODR-10^8^, and its downregulation reveals the much higher mobility of *daf-2* animals^8^, even on food, in addition to its much higher and maintained maximum velocity with age. Previously, we found that *daf-2* worms also extend learning beyond wild-type’s ability^7^, but the full duration of this extension with age was not known. That is, exactly how late in life *daf-2* mutants can still learn and remember, and whether this is proportional to their lifespan extension, was not previously determined.

We previously performed neuron-specific RNA-sequencing in young (Day 1) adult *C. elegans* and identified neuron-specific targets^24^; genes upregulated in *daf-2* mutant neurons are distinct from those in the whole animal, and we found that these neuronal genes are necessary for the observed improvements in memory and axon regeneration in *daf-2* mutant worms. However, whether *daf-2* uses the same or different genes in young and old worms to improve and maintain cognitive function with age is unknown. Recent datasets using whole-animal single-cell RNA-seq have been generated for wild-type and *daf-2* worms, and these are sufficient for whole-body aging and pseudobulk analyses^25–27^, but we have found that those data are not deep enough to use specifically for in-depth analysis of neurons, which can be difficult to gather from whole animals. Other data are from larval stages and cannot be extrapolated to aging adults^28^. To identify the transcriptional differences in the aging nervous system that might contribute to the loss of neuronal function with age in wild-type worms and the differences responsible for the extended abilities of *daf-2* animals, here we performed RNA sequencing on FACS-isolated neurons of aged (Day 8) wild-type and IIS/FOXO mutants. To further investigate the role of the neuronal IIS/FOXO pathway, we identified genes both upregulated by the IIS/FOXO pathway, and genes that are differentially expressed in *daf-2* mutants with age. We found that *daf-2* differentially-regulated genes in the aged neurons are different from young neurons; in fact, many of these Day 8 *daf-2* vs *daf-16;daf-2* upregulated genes are stress response and proteolysis genes that may promote neuronal function and health. We then used functional assays to assess the contributions of *daf-2*-regulated genes to learning and memory. Our results suggest that *daf-*2’s neuronal targets in older worms are required to maintain neuronal functions with age, suggesting that additional and alternative mechanisms are at work in these aged mutants from their young counterparts.

## Results

### Wild-type neurons lose their neuronal function and identity with age

Previously, we found that cognitive abilities in *C. elegans,* including learning, short-term memory, and long-term memory, all decline with age^7^. Moreover, neuronal morphology and regeneration ability are also impaired with age^18,20–22^. However, how these phenotypes are regulated at the molecular level in aging neurons remains to be systematically characterized. Therefore, we were interested in first identifying gene expression changes with age in wild-type neurons to characterize the normal physiological aging process. Before choosing timepoints to assess neuronal transcriptome changes, we carried out associative learning and short-term associative memory assays^7^ as we have previously described^7,29–31^. Briefly, well-fed worms are starved for one hour, then re-fed while exposed to the neutral odorant butanone for one hour; a choice assay between butanone and control immediately after training tests associative learning, while a choice assay after one hour of recovery on food-only plates tests short-term associative memory^7^. Adult Day 1 worms are fully developed, young, and healthy, while wild-type Day 7-8 worms, although still in their mid-life, have completely lost their learning and short-term memory abilities already by Day 7^7^ (**Fig. 1a**), thus we consider them “aged” for the purposes of understanding loss of cognitive ability. Therefore, we reasoned that a comparison of adult wild-type Day 1 neurons with wild-type neurons that are at least aged Day 7-8 should reveal changes with age that result in loss of cognitive function.

**Figure 1.**
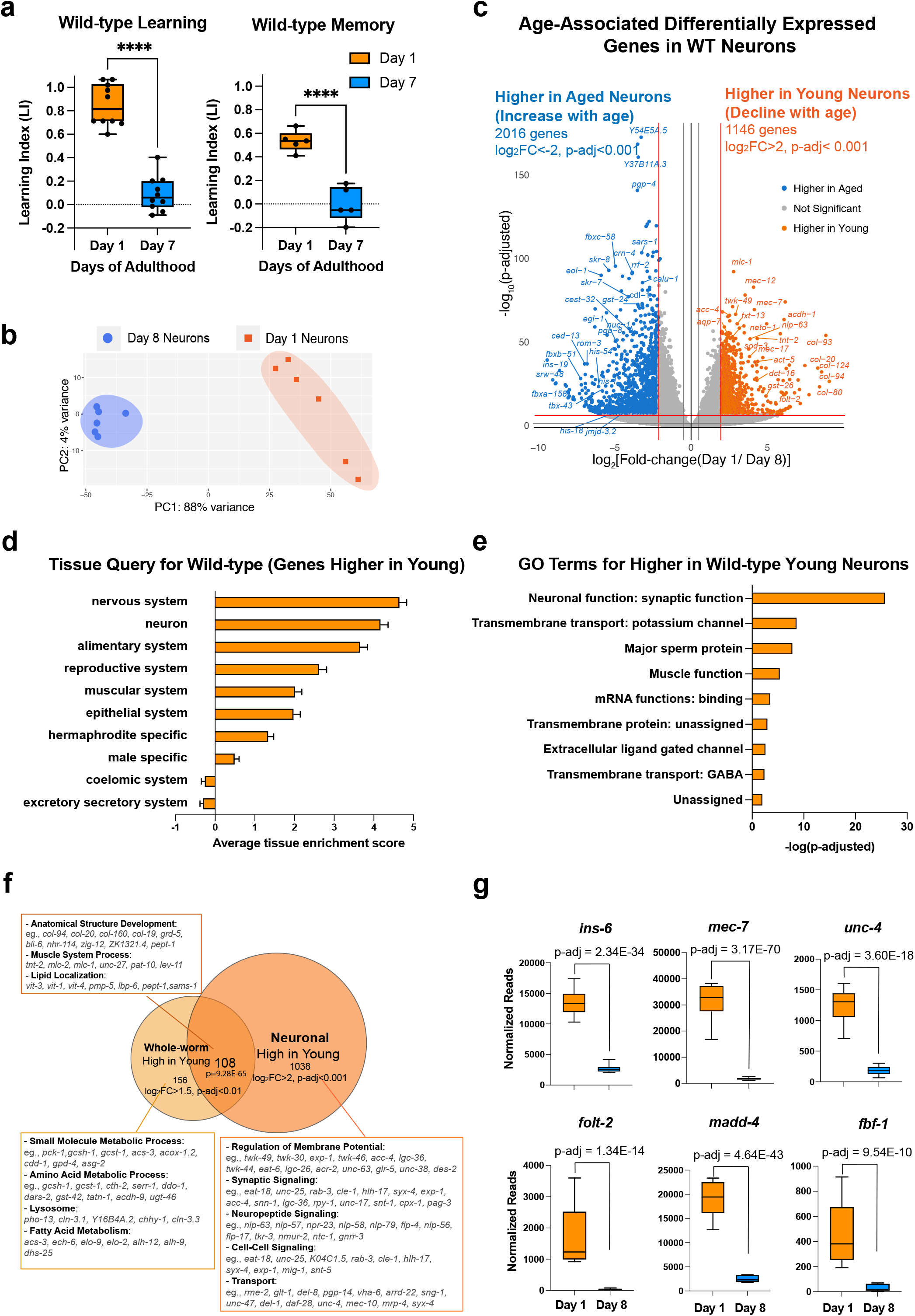
Identifying neuronal aging targets in WT worms using neuron-specific RNA-sequencing. (a) Wild-type learning and 1hr memory results on Day 1 and Day 7. Learning and memory results are represented as learning index (LI). Details of the LI calculation are explained in the methods. Learning, N = 10, memory, N = 5. ****: p < 0.01. Student’s t-test. (b) PCA plot for Day 1 (orange) and Day 8 (blue) neuronal bulk RNA-seq samples. (c) Volcano plot comparing age-associated differentially-expressed genes in WT neurons. Genes downregulated with age (orange) and upregulated with age (blue) were obtained by neuron-specific RNA sequencing of adult wild-type animals with neuron-specific GFP expression. Adjusted p-value < 0.001, log_2_(Fold-change) > 2. N = 6 biological replicates per age. 1146 genes were significantly downregulated with age (higher in young neurons) and 2016 genes were upregulated with age (higher in old neurons) (d) Tissue prediction scores for genes higher in young neurons. (e) GO terms of genes that decline with age in wild-type neurons. Synaptic and signaling GO terms are enriched in neuronal genes. p-value calculated using hypergeometric distribution probability. (f) Comparison of whole-body higher-in-young genes and neuronal higher-in-young genes. GO Terms and representative genes were performed using g:Profiler software. P-value of overlapping regions were calculated using a hypergeometric calculator. (g) Normalized reads of *ins-6, unc-4, mec-7*, *folt-2*, *fbf-1*, and *madd-4*, in Day 1 and Day 8 neurons in our dataset. P-adjusted values were calculated from DESeq2 software. Box plots: center line, median; box range, 25-75^th^ percentiles; whiskers denote minimum-maximum values.

To identify genes that regulate age-related morphological and functional decline in wild-type neurons, we performed neuron-specific transcriptomic analysis using our previous FACS neuronal isolation method^24^ on six biological replicates each of Day 1 and Day 8 adult wild-type worms, where 100,000 GFP+ cells were collected for each sample (**Fig. S1a-c**). Because we previously found that whole-worm analyses mask changes found specifically in neurons^24^, to complement our aging neuron studies, we also carried out RNA-sequencing analyses of aging whole worms (**Fig. S2a-e**), which we found is dominated by changes in the extracellular matrix (**Fig. S2b**), stress response/pathogen genes (**Fig. S2c**) and the alimentary system (intestine) (**Fig. S2d**), overshadowing neuronal changes.

Principal components analysis of the FACS-isolated neuron RNA-seq samples indicated that they are well separated by age (**Fig. 1b**), and downsampling analysis^32^ (**Fig. S1e**) suggested that we have sequenced to saturation, with an average of 41,636,463 uniquely-counted reads and detected the expression of 19725 coding and non-coding genes (log_10_(TPM)>0.5) (**Fig. S1d**). Enrichment analysis of genes that are differentially expressed with age (**Fig. 1c-e**) suggested that neuronal sorting and sequencing were successful because the sequenced genes are enriched for neuronal genes such as *mec-7, mec-12,* and *twk-49*, as expected, and less enriched for all other major tissues. Tissue enrichment analysis of differentially-expressed genes suggested that aging neurons lose genes most expressed in the nervous system and neurons (**Fig. 1d**), as one might expect. Gene ontology (GO) analysis suggested that genes declining with age in neurons encode proteins important in neuronal function (**Fig. 1e, f**), including synaptic proteins (e.g., *srh-59, rab-3, sng-1, sup-1*), potassium channels (e.g., *egl-23*, *twk-7*, *twk-49*, *ncs-5*), and transmembrane transporters (e.g., *folt-2, ccb-2, unc-79, exp-1*). The decrease in expression of these genes during aging may indicate that neurons are losing their identity and their ability to perform neuronal functions, such as signal transduction and axonal transport, and correlates with the behavioral and morphological declines observed in aging wild-type worms.

Comparing whole-worm sequencing and neuron-specific sequencing (**Fig. 1f**), we found that genes involved in metabolic processes decline with age only in the body, and genes encoding structural proteins, lipid localization, and muscle system processes decline with age in both the body and in neurons, while neurons specifically lose genes that are associated with neuronal function, including synaptic proteins, neuropeptide signaling, and other neuron functions, correlating with neuronal loss of function with age. Together, these results indicate that neurons harbor many unique age-related changes that could be overshadowed in the whole-worm transcriptome but are revealed by neuron-specific sequencing.

Many genes that are more highly expressed in young neurons are known to be specific to a subset of neurons. *ins-6*, an insulin-like peptide specific to the ASI, ASJ, and AWA neurons^28^ that regulates longevity^33^ and aversive learning^34^, is significantly downregulated with age (**Fig. 1g**). *srd-23*, a serpentine receptor located at the AWB neuron cilia^35^, also decreases expression with age (**Fig. S1f**). Furthermore, various genes specific to sensory neurons (*txt-12, flp-33*), touch neurons (*mec-7*), and motor neurons (*unc-4*) decline in expression with age (**Fig. 1g, Fig. S1f**). Previous studies showed that loss of genes including *ins-6*^34^, *mec-7*^36^, *unc-4*, *folt-2*^19^, *madd-4*^37^, and *fbf-1*^31^ lead to behavioral dysfunction in motility and chemosensory abilities; therefore, the decreased expression of these neuron type-specific genes with age may impact the function of individual neurons and disrupt neural circuit communication, ultimately contributing to the declines in behavior observed during aging (**Fig. 1g**).

As neurons age, genes that increase in expression, while assigned to the nervous system (**Fig. 2a**) are not specific for neuron function; instead, aged wild-type neurons express higher levels of many predicted F-box genes with predicted proteasome E3 activity (e.g., F-box proteins *fbxa-158*, *fbxb-51*, *pes-2.1*, and SKp1-related proteins *skr-12*, *skr-6*). Some transcription regulation (e.g., *ced-13, tbx-43, nhr-221, end-1*), and chromatin structure and function (e.g., *his-54, dot-1.2, jmjd-3.2, hil-7, utx-1*) genes also increase with age (**Fig. 2b**), even though neurons appear to lose their neuron-specific transcriptional identity with age.

**Figure 2.**
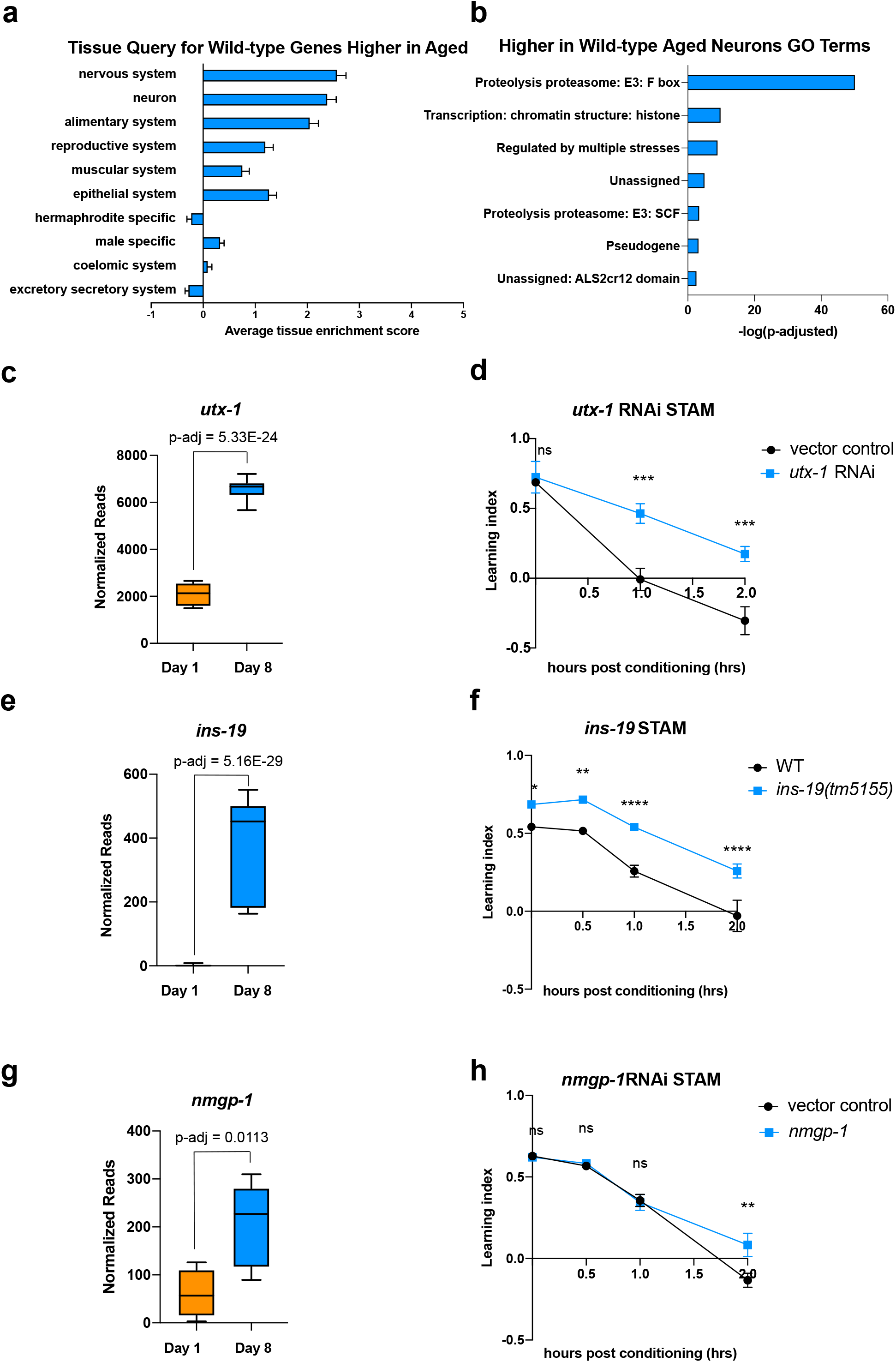
Genes that increase with age cause behavioral defects. (a) Tissue prediction score for wild-type genes expressed at higher levels in aged worms. (b) GO terms of genes expressed higher in aged neurons highlight transcription regulation and proteolysis. GO term analysis was done using Wormcat 2.0. (c) Normalized reads of *utx-1* on Day 1 and Day 8. (d) Short-term associative memory (STAM) assay shows that neuron-sensitized adult-only *utx-1* knockdown improves 1hr and 2hr memory of wild-type worms on Day 2. RNAi was performed using the neuron-RNAi sensitized strain LC108. (e) Normalized reads of *ins-19* on Day 1 and Day 8. (f) *ins-19* mutation improves learning and memory in STAM on Day 3 of adulthood. (g) Normalized reads of *nmgp-1* on Day 1 and Day 8. (h) *nmgp-1* neuron-sensitized RNAi knockdown improves memory in STAM on Day 2. RNAi was performed using the neuron-RNAi sensitized strain LC108.P-adj value of normalized count change generated from DEseq2 analysis. (c,e,g) Box plots: center line, median; box range, 25-75^th^ percentiles; whiskers denote minimum-maximum values. Normalized reads and adjusted p value calculated using the DESeq2 software. Each dot represents one sequencing replicate. (d,f,h) N = 5 plates in each behavioral experiment. Representative result of 2 biological repeats is shown. *: p<0.05. **: p<0.01. ***: p<0.001. ****: p<0.0001. Two-way ANOVA with Tukey’s post-hoc analysis.

One ongoing discussion about changes during aging is how to interpret an increase in expression with age. There are two main models for genes that increase their expression with age and have a resulting impact on function: that they rise with age to compensate for lost function (“compensatory”) and therefore promote function, or that their expression is deleterious to function and only rises with age through dysregulation. If a gene is compensatory, then its knockdown would abrogate learning and memory, even in young animals. If a gene’s function is harmful to neurons, reducing its expression might be beneficial to the worm, even in young animals. (Of course, there may be other scenarios in which a gene with multiple functions may be detrimental for some behaviors but beneficial for other physiological roles.) To test this hypothesis, we reduced expression of a small set of highly-upregulated candidate genes in categories that might function in a compensatory manner. These include *utx-1*, a histone demethylase known to play a role in development^38^ and lifespan in worms^39–41^, and whose homolog has been implicated in cognition in mammals^42,43^; *ins-19,* an insulin-like peptide; and *nmgp-1*, a neuronal glycoprotein involved in chemosensation^44^. In each case, we see that gene expression is significantly higher in old than in young neurons (**Fig. 2c, e, g**). If a gene increases expression to benefit neurons, we would expect to see no difference in memory in young animals where there is no defect; by contrast, if the increase of a gene is deleterious, we would expect to see an improvement in behavior when knocked down, even in young animals. We performed adult-only neuron-sensitized RNAi knockdown to prevent any possible deleterious effects caused by changes during development, which largely takes place in early larval stages; testing in young adult animals is logical because there is no memory in aged wild-type, so any deleterious effect of knocking down a potentially compensatory gene in an aged would not result in a change. For all behavioral assays, we first prioritized significantly-changed genes with high fold-change, and then those with mammalian homologs.

We found that 48 hrs (L4-Day 2) of adult-only knockdown of *utx-1* increases 1hr and 2hr memory (**Fig. 2d**), the loss-of-function mutation of *ins-19* increases both learning and memory (**Fig. 2f**) and the adult-only knock-down of *nmgp-1* extends memory at 2 hours (**Fig. 2h**). That is, in each of these cases, reduction of these genes did not impair memory, as loss of a compensatory function would appear; rather, loss of these age-upregulated genes improved wild-type memory. These results indicate that at least some neuronal genes that increase with age can have a negative impact on learning and memory, as demonstrated by the improvement of memory when knocked down, even in young animals. While it is still possible that some upregulated genes may act in a compensatory manner, the simplest model is that at least some are actively deleterious for learning and memory. We previously observed that for genes that play a role in complex behaviors like learning and memory, the loss of single genes can have a large impact on these complex behaviors^60^, unlike the additive roles of longevity-promoting genes ^11^. Therefore, one mechanism by which wild-type worms lose their learning and memory functions with age is not just by loss of neuronal gene expression, as one might expect, but also by dysregulation of expression of genes that can negatively impact learning and memory.

### *daf-2* mutants maintain learning and memory with age

We previously found that *daf-2* animals have extended motility (Maximum Velocity) that correlates with and predicts their extension of lifespan^8^. Additionally, not only do young *daf-2* worms have better memory than wild-type worms, but *daf-2* mutants also maintain learning and memory better with age^7,24^. However, the duration of this improvement was unknown. To determine the proportion of life that worms can learn and remember, we tested wild-type, *daf-2*, and *daf-16;daf-2* worms for their learning and associative memory ability every day until these functions were lost. We found that while wild-type worms lose their learning and short-term memory abilities by Day 7-8 (**Fig. 1a**, **Fig. 3a, b**), learning and memory span were significantly extended in *daf-2* mutants (**Fig. 3a, b**); thus, a comparison of *daf-2* neurons with wild-type neurons at Day 8 should reveal differences relevant to cognitive aging. The extension of learning and memory is dependent on the FOXO transcription factor DAF-16 (**Fig. 3a**); in fact, while *daf-16;daf-2* mutants still have the ability to learn for a few days, these mutants are completely unable to carry out any memory ability, even on Day 1. Thus, learning ability, which is similar in wild-type and *daf-2;daf-16* mutants, is mechanistically distinct from short-term memory ability^31^. *daf-2* worms maintained learning ability until Day 19 and short-term (1hr) memory ability until Day 15, more than twice the duration of wild-type worms, while *daf-16;daf-2* worms exhibit no short-term memory ability, even on Day 1 of adulthood (**Fig. 3b**). Our data suggest that the learning span-to-lifespan (**Fig. S3c**) and memory span-to-lifespan ratios in *daf-2* worms were similar to or slightly higher than that of wild-type worms (**Fig. 3c, d**), indicating that *daf-2* mutants maintain cognitive function for at least proportionally as long as wild-type worms do. Thus, *daf-2* mutants maintain their higher cognitive quality of life longer than wild-type worms, while *daf-16;daf-2* mutants spend their whole lives without memory ability (**Fig. 3d**), in contrast to claims that *daf-2* mutants are less healthy than wild-type or *daf-16* worms^23^. Additionally it should be noted that because our choice assays distinguish motility function from learning and memory function^29^, the improvements in memory with age shown by *daf-2* mutants relative to wild type are distinct from *daf-2’s* improvements in motility that we previously showed^8^. Therefore, we are interested in these genes that might contribute to the extended cognitive function that *daf-2* worms demonstrate.

**Figure 3.**
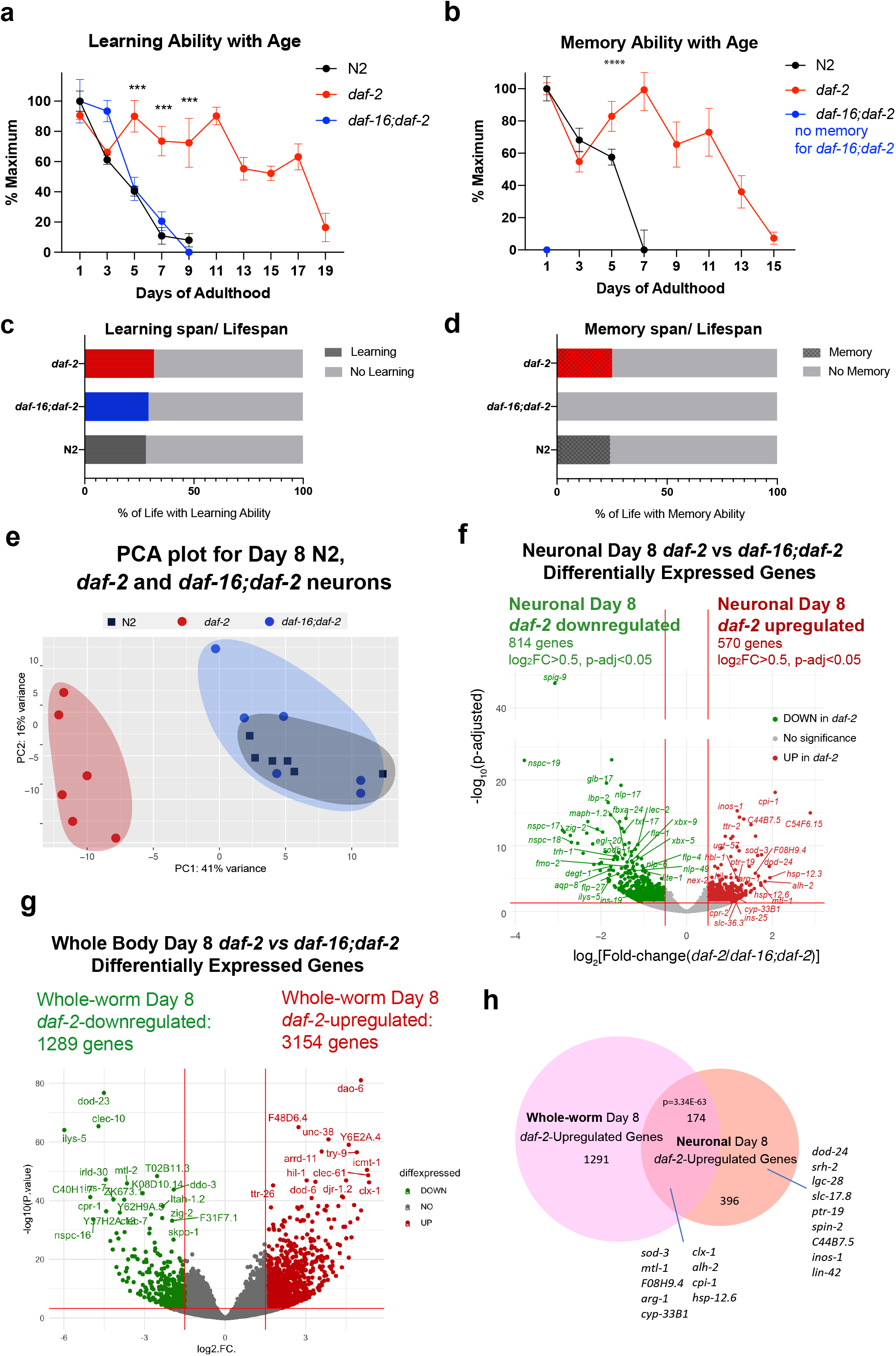
Identifying neuronal IIS/FOXO targets in aged worms using neuron-specific RNA-sequencing. (a) *daf-2* mutants show better learning maintenance with age compared to N2 and *daf-16;daf-2* worms. N = 10 plates in each condition. (b) *daf-2* mutants show better memory maintenance with age compared to N2 worms. *daf-16;daf-2* worms do not have 1hr memory on Day 1 of adulthood. N = 10 plates in each condition. (c-d) *daf-2* mutants have a slightly larger learning span/lifespan ratio and memory span/lifespan ratio than N2 (wild type). Lifespan shown in Figure S3c. (e) PCA plot of Day 8 N2, *daf-2,* and *daf-16;daf-2* neuronal RNA sequencing results. (f) Volcano plot of neuronal *daf-2*-regulated, *daf-16*-dependent up- and downregulated genes on adult Day 8 (Adjusted p-value < 0.05, log_2_(Fold-change) > 0.5, N = 6 biological replicates per strain). 570 genes were significantly upregulated and 814 genes were downregulated in *daf-2* neurons compared with *daf-16;daf-2*. (g) Volcano plot of whole-worm *daf-2* vs *daf-16;daf-2* differentially-expressed genes during aging. 3154 genes are higher in *daf-2*, 1289 genes are higher in daf-*16;daf-2* (log_2_[Fold-change(*daf-2* vs *daf-16;daf-2*)] >1.5, p-adjusted <0.01). (h) Comparison of neuronal and whole-worm Day 8 *daf-2* differentially-expressed genes (overlap p = 3.34E-63, hypergeometric test). Neuron-specific and shared *daf-2* upregulated genes with the highest fold-changes are labeled.

### Aging IIS/FOXO neurons express stress-resistance genes to maintain neuronal function with age

To identify genes that may improve memory and slow cognitive aging in long-lived *daf-2* mutants, we compared the transcriptional profiles of Day 8 FACS-isolated neurons from *daf-2* animals with Day 8 FACS-isolated wild-type and *daf-16;daf-2* neurons; by Day 8, wild-type and *daf-16;daf-2* worms have already lost their learning and memory ability, but *daf-2* worms still maintain their cognitive functions (**Fig. 3a, b**). It should be noted that wild-type worms still have normal chemotaxis and motility at Day 8^7^, and there is a separation of several days between the loss of cognitive functions and the loss of motility^7^; therefore, comparison of the neuronal transcriptomes of *daf-2* with wild type and *daf-16;daf-2* at this age should specifically highlight genes that are required for learning and memory rather than other functions.

The PCA of the *daf-2*, *daf-16;daf-2*, and wild-type neuronal Day 8 transcriptomes (**Fig. 3e**) indicates that aged *daf-16;daf-2* mutant neurons are similar to aged wild-type neurons, correlating well with their similarly worsened cognitive functions at this age; that is, at a transcriptomic level, aged (Day 8) wild-type neurons and aged *daf-16;daf-2* neurons are similar, which is echoed by their shared inability to carry out learning and memory by Day 8 of adulthood (Fig. 3ea, b). By contrast, the transcriptomes of aged *daf-2* mutant neurons are distinct from both aged wild-type and aged *daf-16;daf-2* neuron transcriptomes, just as the cognitive abilities of *daf-2* are much greater than wild type or *daf-16;daf-2* at this age. Downsampling analysis shows that our sequencing depth is sufficient to saturate the detectable differential expression (**Fig. S3g, h**). We obtained an average of 47,233,119 counted reads per sample (Supplemental Table 5) and detected the expression of 16,488 coding and non-coding genes (**Fig. S3e**).

We identified 570 upregulated and 814 downregulated genes in Day 8 *daf-2* neurons compared to Day 8 *daf-16;daf-2* neurons (**Fig. 3f**). A large fraction of the downregulated genes in Day 8 *daf-2* vs *daf-16;daf-2* neurons are “nematode-specific peptide family” (*nspc*-) genes of unknown function (**Fig. 3f)**. While the *daf-2* vs *daf-16;daf-2* changes in whole worms largely replicated the results from our previous studies of young animals^11,45^ (**Fig. 3g, Fig S6g**), comparison of the *daf-2* vs *daf-16;daf-2* differential transcriptional changes in Day 8 whole worms and Day 8 neurons reveal shared (174 genes) and neuron-specific gene expression changes (396 genes; *dod-24, srh-2, lin-42*, etc.) (**Fig. 3h, Fig. S4c**). Not surprisingly, previously-identified genes from whole-worm *daf-2* vs *daf-16;daf-2* and N2 (e.g., *sod-3*, *mtl-1, cpi-1, hsp-12.6,* etc.) that play roles in both neurons and other tissues even in Day 1 *daf-2* mutants appear in the shared list. (Some neuron-specific Day 8 *daf-2-*upregulated genes have not been reported to be expressed in neurons previously (e.g., *spin-2*), further suggesting the value of transcriptomic analyses of isolated neurons in mutant backgrounds at this age.)

Many genes upregulated in Day 8 *daf-2* neurons relative to *daf-16;daf-2* are related to stress responses, including heat stress (e.g., *hsp-12.6, hsp-12.3, F08H9.4/hsp*), oxidative stress (e.g., *sod-3*), and metal stress genes (e.g., *mtl-1*); and proteolysis (e.g., *cpi-1*, *cpr-2*, and *tep-1*). The upregulation of these genes may perform neuroprotective functions, as their homologs in mammals have been shown to do (**Table 1**). Specifically, 36 of the top 100 upregulated genes have identified orthologs or identified domains with known function, of which 32 of (89%) have functions in promoting neuronal health. These mammalian homologs protect neurons against protein aggregation and harmful metabolites (e.g., *cpi-1, alh-2, ttr-41, gpx-5*)^46–50^, maintain synaptic organization and neuronal homeostasis (e.g., *dod-24, ptr-19, plep-1*)^51–53^, facilitate neuronal injury repair (e.g., *F08H9.4, sod-3*)^54,55^, and maintain normal neuronal function (e.g., *lgc-28, slc-36.3, lin-42*)^56–59^. Together, these genes may help maintain *daf-2*’s neuronal health and protect neurons from accumulation of environmental harm during aging.

We found that about a third of the *daf-2*-upregulated genes were shared between the *daf-2* vs *daf-16;daf-2* analysis and the *daf-2* vs N2 analysis (338 genes) (**Fig. S5**). Of the unshared genes, the *daf-2*-maintained genes that are specific to the *daf-2* vs N2 comparison are bZIP transcription factors, including *zip-5, zip-4, atf-2*, and proteasome components (Figure S6D). These results indicate that other transcription factors may participate in regulating *daf-2* functions in aged neurons in addition to the *daf-16*/FOXO transcription factor.

### IIS/FOXO transcriptomic changes are necessary for *daf-2* mutant’s improved neuronal functions

We were interested not only in the genes that remained upregulated with age, but also in genes that might have increased with age in the high-performing *daf-2* mutants. That is, are there genes that increase in expression in *daf-2* mutants that are necessary or beneficial for their continued high performance with age? Some of the Day 8 *daf-2* vs wild-type or *daf-16;daf-2* upregulated genes are also Class 1 DAF-16-dependent genes^11^ (*sod-3, hsp-12.3, fat-5,* and *mtl-1, hil-1*, and *dao-2*). However, many more genes were differentially expressed in Day 8 *daf-2* vs *daf-16;daf-2* neurons from our Day 1 data^24^ (**Fig. 4a, Fig. S4d**). Of the “new” genes – that is, genes upregulated specifically in neurons of Day 8 vs Day 1 of *daf-2* vs *daf-16;daf-2* – many have mammalian homologs that have been shown to play neuroprotective roles, by protecting against aggregation proteins and harmful metabolites, maintaining synaptic organization, neuronal homoeostasis, or neuronal activity, or facilitating neuronal injury repair (see **Table 1** for specific references).

**Figure 4.**
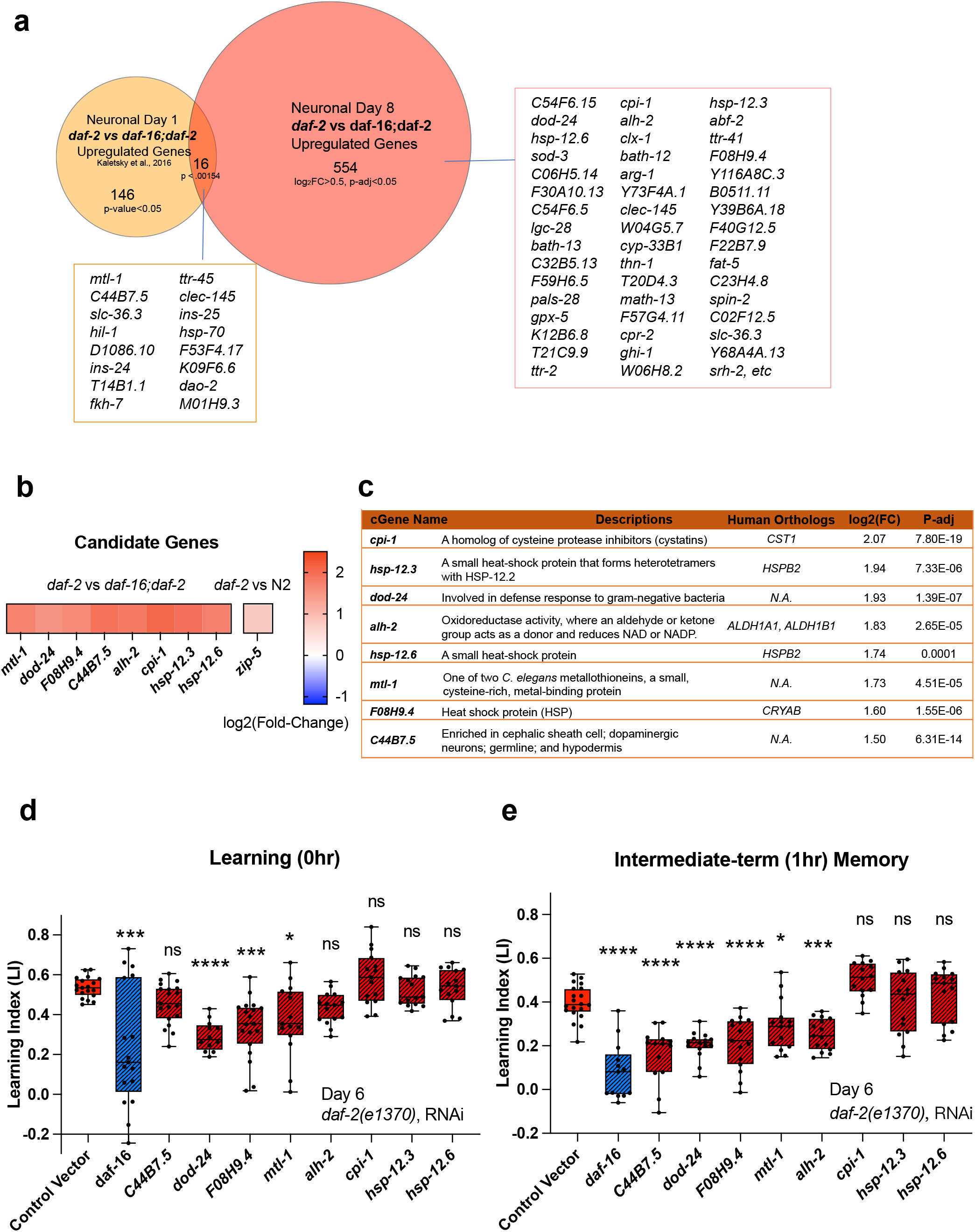
Neuronal IIS/FOXO aging targets regulate memory decline with age in *daf-2* worms. (a) Comparison of neuronal Day 1 and Day 8 *daf-2* vs *daf-16;daf-2* upregulated genes. All shared genes and top Day 8-specific *daf-2* upregulated genes are labeled. (b) *daf-2*-regulated fold-change profile of candidate genes. All candidates are upregulated in *daf-2* mutants. (c) Description of candidate genes. log_2_(Fold-change) and p-adjusted values from the *daf-2* vs *daf-16;daf-2* comparison unless stated otherwise. (d) Candidate gene knockdown effects on Day 6 adult *daf-2* learning (0hr after conditioning). Two candidate genes, *dod-24* and *F08H9.4*, show a significant decrease in learning ability. N = 5 plates in each condition, merged results of 3 biological repeats shown. (e) Candidate gene knockdown effects on Day 6 adult *daf-2* short-term memory (1hr after conditioning). *C44B7.5*, *dod-24*, *F08H9.4*, *mtl-1,* and *alh-2* showed significant decreases in memory. N = 5 plates in each condition, the representative image of 3 biological repeats shown. (d-e) RNAi was performed using a neuron-sensitized RNAi strain CQ745: *daf-2(e1370) III;* vIs69 [*pCFJ90(Pmyo-2::mCherry + Punc-119::sid-1)*] *V*.*: p < 0.05. **: p < 0.01. ***: p<0.001. ****: p <0.0001. One-way ANOVA with Dunnet’s post-hoc analysis. Box plots: center line, median; box range, 25-75^th^ percentiles; whiskers denote minimum-maximum values.

If the upregulated genes in aged *daf-2* neurons are responsible for the extended memory span of *daf-2* mutants, knocking down those genes should block older *daf-2* mutants’ memory functions. Therefore, we tested the effect of RNAi knockdown of the top fold-change candidate genes on *daf-2’s* memory in aged adults. We chose Day 6 for testing because by then, like on Day 8, wild-type worms have already lost their learning and most memory abilities, but *daf-2* worms retain normal cognitive functions, and this time point avoids the increased naïve chemotaxis that we observe in older *daf-2* animals. As shown in **Fig. 4b-c**, we selected these significantly differentially-expressed candidate genes based on their ranking in fold-change. Previously, we have found that the top significantly differentially-expressed genes (by fold-change) are most likely to have strong effects on function, while less differentially-changed genes have less of an effect^11,24,60^, therefore we prioritized genes that are significantly different and the most highly expressed in *daf-2* mutants compared to *daf-16;daf-2* mutants for subsequent testing (**Fig. 4c**). *daf-2* worms, including neurons, are susceptible to RNA interference^24,61^. Of the eight candidate genes we tested, the reduction of three of them (*F08H9.4, mtl-1,* and *dod-24*, originally classified as a Class II gene with proposed immune activity) significantly decreased *daf-2’s* learning ability on Day 6 (**Fig. 4d**). Those genes plus reduction of two additional genes (*C44B7.5* and *alh-2*) affected 1hr memory (**Fig. 4e**) in Day 6 *daf-2* mutants. That is, knockdown of the heat shock-related gene *F08H9.4*, the innate immunity gene *dod-24*, aldehyde dehydrogenase *alh-2* and previously uncharacterized gene *C44B7.5* are required to some degree for *daf-2’s* extended memory ability. The reduction of the metal stress gene *mtl-1*, which is expressed in neurons as well as the rest of the body, had a slight effect on learning and memory.

One caveat of these experiments is that, while we found these genes through the isolation of neurons from aged worms and subsequent RNA-sequencing, the knockdown of the genes and its effects are not necessarily neuron-autonomous; however, *alh-2* and *F08H9.4* were reported to only be expressed in neurons and the cephalic sheath cell^62^, and *C44B7.5* and *dod-24*, while expressed more broadly, were not upregulated in *daf-2* in the whole-worm analysis (Fig. 3f), therefore their effects are most likely neuron-autonomous. In fact, *dod-24* is one of the original Class 2 *daf-2*-downregulated genes from whole-animal analyses, suggesting that *dod-24’s* increase in expression is specifically in neurons, therefore the effect of its knockdown is most likely to be neuron-autonomous.

Together, these data suggest that the specific genes that are differentially regulated in Day 8 *daf-2* mutants may aid in slowing neuronal function decline and behavioral changes associated with aging. Furthermore, memory maintenance with age might require additional genes that function in promoting stress resistance and neuronal resilience, which were not previously uncovered in analyses of young animals.

## Discussion

Although it has been shown previously that *daf-2* worms maintain various functions with age, how long they can maintain learning and memory with age, and the genes that might be responsible for these extended neuronal functions, have not been previously explored. Here we have found that *daf-2* worms maintain learning and memory abilities proportional with (or even slightly beyond) their degree of lifespan extension, underscoring *daf-2’s* improved healthspan^8^. To understand how memory is lost with age and retained in insulin/IGF-1-like signaling mutants, we have characterized the neuronal transcriptomes of aged wild-type worms and IIS (*daf-2*) and IIS/FOXO (*daf-16;daf-2*) mutants (**Fig. 5**). We found that wild-type neuronal aging is characterized by a downregulation of neuronal function genes and an upregulation of proteolysis genes and transcriptional and epigenetic regulators, which together may help explain the loss of neuronal identity and function with age. We also identified the transcriptomic profile accompanying *daf-2*’s extended learning and memory span. Specifically, *daf-2* neurons maintain higher expression of stress response genes and predicted neuronal homeostasis functions (**Table 1**), which may help make them more resistant to environmental adversities and age-related decline. We also identified genes responsible for wild-type worms’ worsened learning and memory with age.

**Figure 5.**
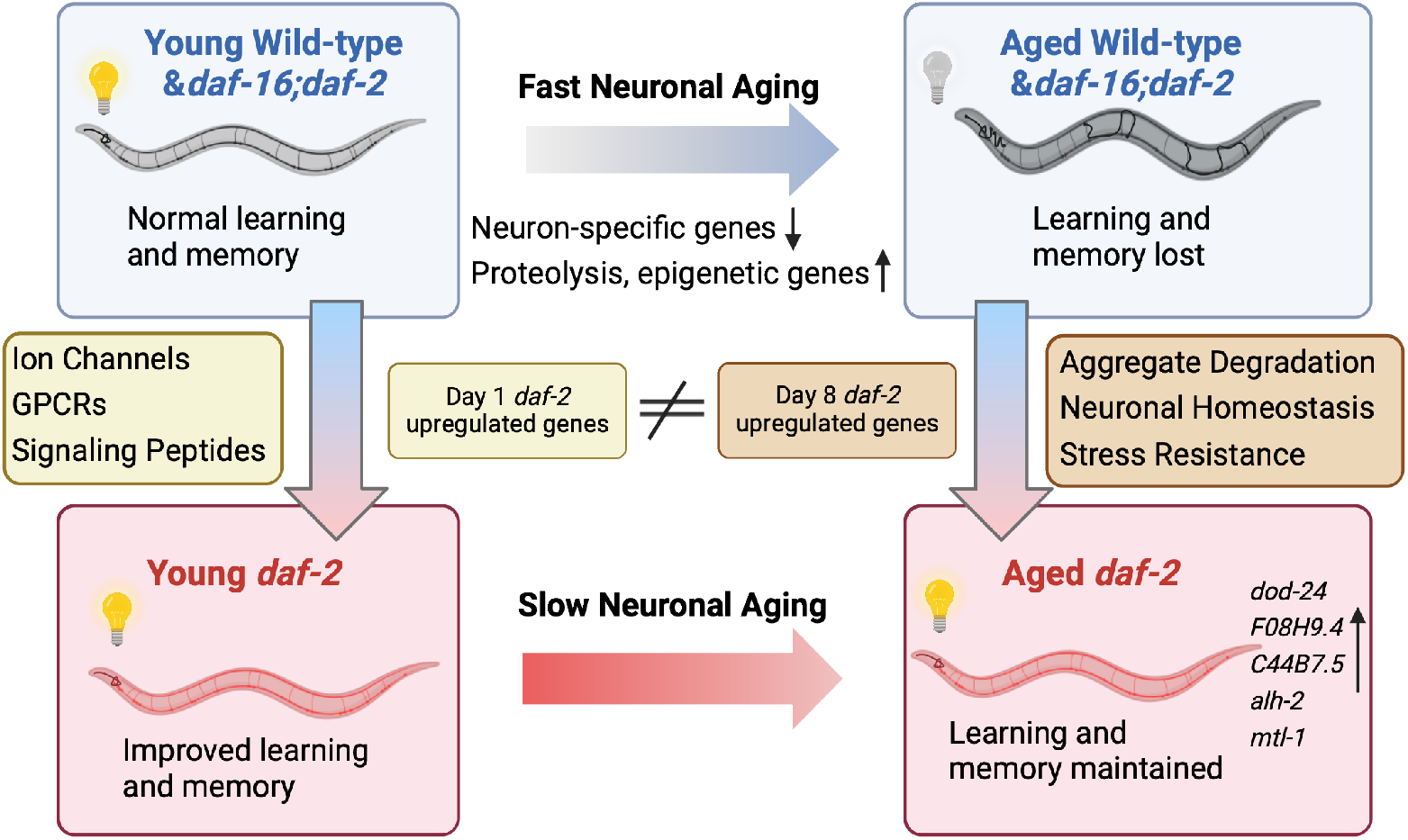
Aged *daf-2* neurons upregulate neuroprotective genes to maintain improved cognitive behaviors. During normal neuronal aging, neuron-specific genes decrease in expression, while proteolysis and epigenetic regulators are upregulated, resulting in neuron dysfunction and cognitive function loss. In aged *daf-2* mutants, upregulation of neuroprotective genes including *dod-24*, *F08H9.4*, *C44B7.5*, *alh-2*, and *mtl-1* contribute to *daf-2*’s improved cognitive function. The diagram was generated using Biorender.

By employing a FACS-based neuron-sorting technique, we selectively analyzed adult neuron-function-related genes and investigated their aging process, which is not easily discernible through whole-worm sequencing^25–27^ (**Fig. S6**). Sequencing many biological repeats of aging neurons to high depth with ribosomal RNA depletion allowed us to detect a larger number of genes compared to other neuron-related bulk and single-cell sequencing profiles^26^, providing a deep transcriptomic dataset of aged wild-type, IIS mutant, and IIS/FOXO mutant neurons. Our analysis allowed us to identify differentially-expressed genes that are known to be expressed in at a small number of neurons, even for low-abundance genes. Notably, our sequencing results uncovered genes previously not known to be expressed in neurons that remained undetected in other datasets. Moreover, we revealed the involvement of known neuronal genes in the aging process, such as *ins-6* and *srd-23*. We hope that this dataset will become a valuable resource for detecting new candidates in neuronal aging.

For example, *dod-24*, which we observed to be upregulated in *daf-2* neurons and required for *daf-2*’s extended memory, was downregulated in the *daf-2* whole-worm transcriptome (**Fig. S4c**). *dod-24* has been traditionally classified as a Class II gene that is downregulated in *daf-2* worms and upregulated by *daf-16* RNAi treatment^11,45^. Functionally, it has been shown to be an innate immunity gene upregulated during pathogen infection^63–65^, and its whole-body reduction has been shown to extend the lifespan of wild-type animals^11^. However, here we find that *dod-24* is beneficial in the nervous system and required for *daf-2*’s extended learning and memory in aged worms. This intriguing contrast between the whole-worm transcriptome and the neuron-specific transcriptome suggests that some genes may have distinct regulatory roles in the nervous system, necessitating a more precise approach beyond whole-worm transcriptomics.

Using this neuron-specific sequencing profile of aged cells, we identified key pathways that change during neuron aging. Our sequencing of aged neurons uncovered active transcriptomic alterations during aging, resulting in not just transcriptional silencing but also upregulation of various pathways. We found that the Day 8 *daf-2* vs *daf-16;daf-2* neuronal differentially-expressed genes that we have newly discovered here differ from the neuronal Day 1 *daf-2* vs *daf-16;daf-2* dataset we previously obtained^24^. These Day 8 differentially-expressed genes are not canonical neuronal genes, such as receptors and ion channels; instead, there are more metabolic and proteolytic genes whose protein orthologs have been shown to be neuroprotective. All top 50 genes and 90% of the top 100 genes with identified mammalian orthologs have been shown to be essential to neuronal functions in mammals (Table 1). These results indicate that instead of simply mimicking a young state, *daf-2* may enhance neuron’s resilience to accumulation of harm and take a more active approach to combat aging. These changes suggest that *daf-2*’s extended memory maintenance may require different mechanisms than function in young animals; *daf-2* may maintain neuronal function not just by retaining a youthful transcriptome, but also by increasing the expression of genes that promote resilience, such as stress-response genes and proteolysis inhibitors.

In addition to examining aging in wild-type and IIS/FOXO mutants independently, our results further linked the normal aging process to altered gene regulation in the IIS pathway. *utx-1, nmgp-1,* and *ins-19* increase in expression in aged neurons, and we found that their reduction improved memory, indicating that at least some of the genes whose expression rises with age can have a negative impact on normal cognitive functions, rather than acting in a compensatory manner. *utx-1* is an H3K27me3 histone demethylase we found to be higher in wild-type aged neurons, but it is also involved in the IIS pathway. The downregulation of *utx-1* has been shown to regulate development^38^ and promote longevity^39–41^, and its mammalian homolog has been implicated in regulating cognitive abilities^42,43^. The longevity response of *utx-1* depends on *daf-16*^39–41^. The loss of *utx-1* decreases methylation on the *daf-2* gene, thus increasing DAF-16’s nuclear localization, mimicking a *daf-2* mutation^39^. This example of the crosstalk between normal aging and IIS/FOXO mutants offers valuable insights into modifying the aging process for enhanced longevity and cognitive health.

We found that the insulin-like peptide *ins-19* was upregulated in aged neurons and was downregulated in aged *daf-2* neurons, and its downregulation in wild-type worms extended memory span. Insulin-like peptides play crucial roles as receptor ligands (in both agonist and antagonist roles) for DAF-2, and we have found them to be downregulated in *daf-2* mutants compared with *daf-16;daf-2* mutants, possibly creating a feedback loop that dampens the insulin signaling pathway, as was previously shown for *ins-7* and *ins-18*^11,66^. These peptides exhibit diverse functions in development, dauer formation, and longevity^11,66–70^. Notably, certain insulin-like peptides have been linked to neuronal activities, such as the regulation of aversive learning by the two antagonistic peptides *ins-6* and *ins-7*^34^, and reduced long-term learning and memory by *ins-22* RNAi^60^. In our study, the expression changes of *ins-19* during wild-type aging and in *daf-2* mutants provide an example of how longevity mutants can reverse wild-type transcriptional changes during aging, ultimately reducing behavioral and functional decline.

### Summary of mechanistic insights

Our analysis of transcriptomes from isolated aged Day 8 neurons of wild-type, *daf-2,* and *daf-16;daf-16* mutants reveal several major mechanistic insights. Specifically, we found that wild-type neurons lose their neuronal identity through a combination of the loss of neuron-specific function genes with age, and the concomitant dysregulated increase in non-neuronal genes with age. Furthermore, at least a fraction of the top-upregulated genes with age can play deleterious roles; that is, they rise with age, and their knockdown improves function, even in young animals. This argues against the idea that all of these genes are playing a compensatory role with age.

We also found that the knockdown of individual top-ranked genes that function in learning and memory can have a large impact - like removing a cog of a machine. This is in contrast to our earlier findings regarding gene reduction in lifespan, where most cellular longevity processes regulated by DAF-16 activity appear to be additive, and therefore loss of individual major genes downstream of DAF-2 and DAF-16 have at most a 5-10% impact^11^. Several of these genes we found to be required for *daf-2*’s age-related improvement in learning and memory - namely *dod-24*, *F08H9.4*, *C44B7.5*, and *alh-2* – were previously not associated with memory function. Finally, these genes are distinct from the set of upregulated Day 1 *daf-2* vs *daf-16;daf-2* genes; how they each individually maintain neuronal function better with age will be interesting to dissect.

### Conclusions

Beyond our sequencing analysis, we have established links between genomics, function, and behavior. We also identified several new genes required for *daf-2*’s age-related improvement in learning and memory, shedding light on their neuron-specific roles. These additional findings further suggest that neuronal sequencing datasets can be used to identify functional candidates and pathways during the aging process. By bridging the gap between transcriptomic landscapes, genetic regulation, and functional outcomes, our study provides a greater understanding of the mechanisms underlying neuronal aging, providing insights into development of aging interventions.

## Methods

### Strains and worm cultivation

N2 (wild type), OH441: otIs45(*unc-119::GFP*), CQ295: otIs45(*unc-119::GFP*);*daf-2(e1370)*, CQ296: otIs45(*unc-119::GFP*); *daf-16(mu86);daf-2(e1370)*, LC108: uIs69 (*myo-2p::mCherry* + *unc-119p::sid-1*), CQ705: daf-2(e1370) III, 3X outcrossed, CQ745: *daf-2(e1370) III;* vIs69 [*pCFJ90(Pmyo-2::mCherry + Punc-119::sid-1)*] *V*, QL188: *ins-19(tm5155) II*, CX3695: kyIs140(*str-2::GFP + lin-15(+)*), CQ461*: (daf-2(e1370);Pmec-4::mCherry)* and CQ501*: (daf-2 (e1370);zip-5(gk646);Pmec-4::mCherry)*. Strains were grown on high-growth media (HGM) plates seeded with E. coli OP50 bacteria using standard methods ^71^.

### Tissue-specific isolation

For neuronal isolation, five plates of fully-grown worms from HG plates were synchronized by hypochlorite treatment, eggs spread on seeded plates to hatch, and at least five plates/replicate were grown to L4 on HGM plates until transferred to HGM plates with FUdR to avoid progeny contamination. This gives us ∼6000 healthy Day 8 worms to sort. Neuron isolation and Fluorescent-activated cell sorting were carried out as previously described ^24,62^. Briefly, worms were treated with 1000uL lysis buffer (200 mM DTT, 0.25% SDS, 20 mM HEPES pH 8.0, 3% sucrose) for 6.5 mins to break the cuticle. Then worms were washed and resuspended in 500uL 20mg/mL pronase from *Streptomyces griseus* (Sigma-Aldrich). Worms were incubated at room temperature with mechanical disruption by pipetting until no whole-worm bodies were seen, and then ice-cold osmolarity adjusted L-15 buffer(Gibco) with 2% Fetal Bovine Serum (Gibco) were added to stop the reaction. Prior to sorting, cell suspensions were filtered using a 5um filter and sorted using a FACSVantage SE w/ DiVa (BD Biosciences; 488nm excitation, 530/30nm bandpass filter for GFP detection). Sorting gates were determined by comparing with age-matched, genotype-matched non-fluorescent cell suspension samples. Fluorescent neuron cells were directly sorted into Trizol LS. 100,000 GFP+ cells were collected for each sample.

### RNA extraction, library generation and sequencing

We used standard trizol-chloroform-isopropanol method to extract RNA, then performed RNA cleanup using RNeasy MinElute Cleanup Kit (Qiagen). RNA quality was assessed using the Agilent Bioanalyzer RNA Pico chip, and bioanalyzer RIN>6.0 samples were observed before library generation. 2ng of RNA was used for library generation using Ovation SoLo RNA-Seq library preparation kit with AnyDeplete Probe Mix-*C. elegans* (Tecan Genomics) according to the manufacturer’s instructions ^72^. Library quality and concentration was assessed using Agilent Bioanalyzer DNA 12000 chip. Samples were multiplexed and sequencing were performed using NovaSeq S1 100nt Flowcell v1.5 (Illumina).

### Data processing

FastQC was performed on each sample for quality control analysis. RNA STAR package was used for mapping paired-end reads to the *C. elegans* genome ce11 (UCSC Feb 2013) using the gene model ws245genes.gtf. Length of the genomic sequence around annotated junctions is chosen as read length −1. 50-70% reads were uniquely mapped. Reads uniquely mapped to the genome were then counted using htseq-count (mode = union). DESeq2 analysis was then used for read normalization and differential expression analysis on counted reads ^73^. Genes with a log_10_TPM >0.5 were considered as detected and genes with a log_2_(fold-change) > 0.5 and p-adjusted <0.05 are considered differentially expressed in further analysis. Gene ontology analysis were performed using gprofiler^74^ or WormCat 2.0^75^ and category 2 was selected to show. Tissue query was performed on top 500 highest fold-change genes, using worm tissue query website (www.worm.princeton.edu)^62^, and only major systems were selected in the analysis.

### Learning and Memory Experiments

We performed Short-Term Associative Memory (STAM) experiments as previously described ^7,29^. Briefly, we used five plates of synchronized adult worms/sample to perform each experiment. One plate of worms was used to test naïve chemotaxis assay without conditioning, while the other 3 plates were washed into M9, and washed 3 additional times to get rid of the bacteria. These worms are starved for 1hr to prime them for food uptake. Then these worms are transferred to conditioning plates with NGM plates seeded with OP50 and 10% butanone stripes on the lid for 1hr to perform conditioning. After conditioning, worms are either transferred from the conditioning plate directly to the chemotaxis plates to assess learning, or transferred to holding plate for 1hr or 2hrs to assess for memory. After staying on holding plates for 1hr or 2hrs, worms are then transferred onto chemotaxis plates to assess for short-term memory. Chemotaxis assays were performed by transferring worms onto chemotaxis plates with 1uL 10% butanone and 1uL ethanol spots separated by 8cm on a 10cm NGM plate. Worms who have reached either butanone spot or the ethanol spot are paralyzed by the 1uL 7.5% NaN3 on these spots. For each timepoint, 5 chemotaxis plates are used to minimize the variation of the outcome. We performed this chemotaxis assay to butanone on naïve and appetitive-trained worms at different time points to assess change in preference to butanone. Chemotaxis index is calculated as (# of worms at butanone-# of worms at ethanol)/(total # of worms - # of worms at origin).

Learning index is calculated by subtracting trained chemotaxis index with naïve chemotaxis index.

For learning and memory span assays, we obtained synchronized worms from hypochlorite-treated eggs. Synchronized worms were washed onto 5’-fluorodeoxyuridine (FUdR) at L4 and maintained on FUdR plates by transferring to new plates every 2 days. 1 Day Prior to experiments, worms are washed onto fresh HG plates without FUdR to avoid change in behavior caused by FUdR. To verify that FUdR has no effect on short-term memory, we compared worms with and without FUdR (**Fig. S6d**), and found no differences. For *utx-1* and *nmgp-1* RNAi experiments, synchronized L4 neuron-RNAi sensitized worms were washed onto HGM plates with carbenicillin and IPTG and seeded with HT115 RNAi bacteria containing the RNAi constructs from the Ahringer Library. For *daf-2* upregulated candidates’ RNAi experiments, synchronized L4 *daf-2* neuron-RNAi-sensitized worms were washed onto HGM plates added with carbenicillin, FUdR, and isopropyl-b-D-thiogalactopyranoside (IPTG) and seeded with HT115 bacteria containing RNAi constructs generated from the Ahringer RNAi Library, then were transferred onto fresh RNAi plates every 2 days until Day 6. 1 Day Prior to experiments, worms are transferred onto plates without FUdR.

### Quantitative and Statistical Analysis

All experimental analysis was performed using Prism 8 software. Two-way ANOVA with Tukey post-hoc tests were used to compare the learning curve between control and experimental groups. One-way ANOVA followed by Dunnet post-hoc tests for multiple comparisons was performed to compare learning or 2hr memory between various treatment groups and control. Chi-square test was performed to compare the neuron morphology change between young and aged AWC neurons. All GO term analysis were perform using Wormcat 2.0 software with Bonferroni corrected adjusted p-values. Venn diagram overlaps were compared using the hypergeometric test. Differential expression analysis of RNA-seq were performed using DESeq2 algorithm and adjusted p-values were generated with Wald test using Benjamini and Hochberg method (BH-adjusted p values). Additional statistical details of experiments, including sample size (with n representing the number of chemotaxis assays performed for behavior, RNA collections for RNA-seq, and the number of worms for microscopy), can be found in the methods and figure legends. Regression analyses were performed using sklearn packages. Correlations were calculated using the SciPy packages.

## Data availability

Sequencing reads are deposited at NCBI BioProject under accession number PRJNA999305.

## Code availability

Analysis codes generated from this study are available upon request.

## Resource availability

Further information and requests for resources and reagents should be directed to and will be fulfilled by Coleen T. Murphy.

## Acknowledgments

We thank the Caenorhabditis Genetics Center (CGC) for strains, WormBase (version WS289) for information, Jasmine Ashraf, William Keyes, Yichen Weng, and Titas Sengupta for help with the experiments, R. Arey and other Murphy lab members for an early replicate of *daf-2* learning and memory with age, members of the Murphy Lab for input on the manuscript, Christina DeCoste, Katherine Rittenbach and the Flow Cytometry Facility for cell sorting assistance, Wei Wang and the Genomics Facility for sequencing assistance, Lance Parsons, Bruce Wang, and Chen Dan for insights on data analysis, and Biorender.com for schematic design. C.T.M. is the Director of the Simons Collaboration on Plasticity in the Aging Brain (SCPAB), which supported the work, and the Glenn Center for Aging Research at Princeton. Y.W. and S.Z. are supported by China Scholarship Council (CSC). K.M. is supported by HHMI Gilliam Fellows Program.

## Author Contributions

C.T.M., R.K., S.Z., and Y.W. conceived and designed experiments. Y.W. and S.Z. performed RNA-sequencing experiments. Y.W., S.Z., and K.M. performed behavioral assays. Y.W. performed imaging experiments. Y.W., S.Z., K.M., S.L., R.K., and C.T.M. analyzed and interpreted results. Y.W. and C.T.M wrote the paper.

---

**Table 1.**
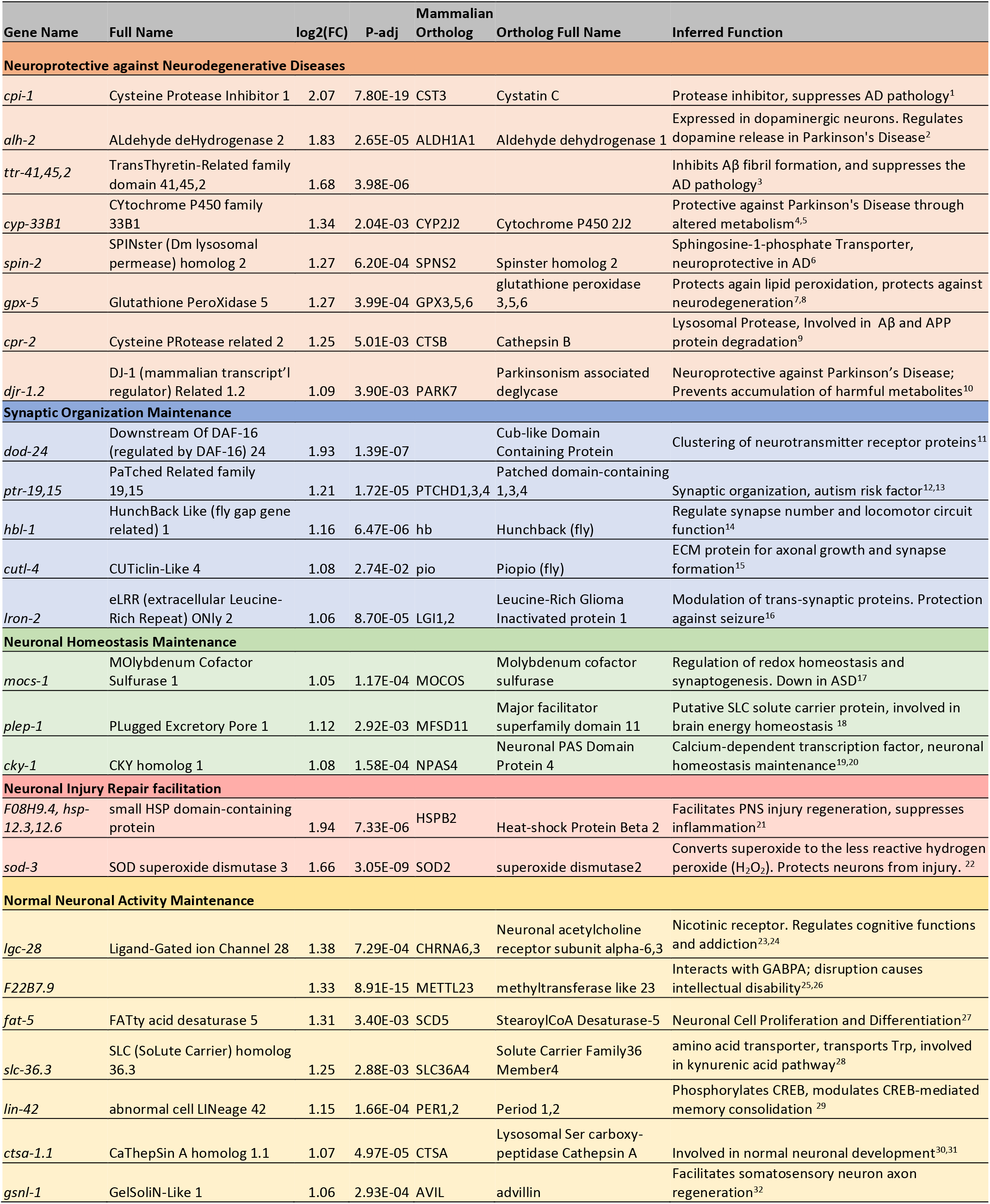
List of top *daf-2* vs *daf-16;daf-2* upregulated genes with orthologs that have neuroprotective functions.

