## Supplementary information for "The Neuron-specific IIS/FOXO Transcriptome in Aged Animals Reveals Regulatory Mechanisms of Cognitive Aging"

**Supplemental Figures**

**Figure S1. Comparison with recent sequencing datasets.** (a) Comparison with CeNGEN (L4 stage) gene expression data[28] shows high correlation, with many genes only detected in our isolated neuron bulk-sequencing dataset (orange; wild-type Day 1 neuron sequencing data used in the comparison). Genes with an average log2(TPM) > 0.5 are considered detected. (b) Venn diagram showing genes detected in this dataset (orange), CeNGEN dataset (brown) and both (gray); only wild-type Day 1 neuron sequencing data were used in the comparison. (c-d) Comparison with gene expression data from the age-matched neuron cells from the Roux et al., dataset[27] shows high correlation, with many genes only detected in our bulk-sequencing dataset. (e-f) Venn diagram showing genes detected in this dataset (orange or blue), Roux et al., dataset (brown) and both (gray). Only age-matched neuron cells from the Roux et al., dataset was used for gene expression analysis. (g) Correlation between our whole-worm Day 8 *daf-2* and *daf-16;daf-2* sequencing and Class I and Class II gene rank from Tepper et al., 2013[45]. High correlation indicates consistency between our isolated neuron RNA-sequencing results with former microarray results, despite the differences in approaches and ages. (h) STAM with and without FUdR shows that 3 Days on FUdR does not affect learning and memory ability.

**Figure S2. Aged neuron-specific sequencing.** (a-b) FACS results of neuron isolation. Over 99.94% of the cells collected are GFP+ neurons. 100,000 cells are collected for each biological replicate, 6 biological replicates for each condition. (c) Workflow of neuron isolation, library generation and sequencing. (d) Number of genes detected in Day 1 and Day 8 wild-type neurons. Genes with $\log_2$(TPM)>0.5 are considered expressed. (e) Down-sampling analysis for N2 Day 1 vs Day 8 indicates downsampling 30% of the data will still yield good results, indicating sufficiency of sequencing depth. (f) Normalized reads of *txt-12, flp-33*, and *srd-23* in Day 1 and Day 8 neurons. P-adjusted values were calculated from DESeq2 software.

**Figure S3. Whole-worm RNA-sequencing identifies whole-body changes during aging.** (a) Volcano plot of Day 1 vs Day 8 differentially-expressed genes during aging. 264 genes are expressed at higher levels in young worms, 1626 genes are higher in aged worms ($\log_2$[Fold-change (Day 1/Day 8)] >2.0, p-adjusted <0.001). (b) GO terms of genes that are expressed at higher levels in young (wild-type) whole animals highlight collagen and metabolism. (c) GO terms of genes that are expressed at higher levels in aged (wild-type) whole animals. GO terms were generated using Wormcat 2.0. (d) Tissue query for whole-worm aged-related genes highlights the alimentary system. (e) Full image of comparison of top wild-type neuronal and whole-worm differentially expressed with age genes. Related to Figure 2D.

**Figure S4. Neuron-specific sequencing of Day 8 *daf-2* and *daf-16;daf-2* mutants.** (a-b) FACS results of neuron isolation. Over 99% of the cells collected are GFP+ neurons. 100,000 cells are collected for each biological replicate, 6 replicates for each genotype. (c) Representative image of N2 and *daf-2* lifespan. (d) Number graph of lifespan, learning, and memory function. Related to Figure 4C and 4D. (e) Number of genes detected in Day 8 N2, *daf-2,* and *daf-16;daf-2* neurons. (f) Ribosomal RNA depletion during sequencing. We used the library generation protocol with *C. elegans*-specific ribosomal RNA depletion kit (Tecan Genomics) successfully depleted rRNA to less than 20% of total reads. (g-h) Downsampling analysis for daf-2 vs *daf-16;daf-2* and daf-2 vs N2. Both shows sufficient depth.

**Figure S5. Whole-worm RNA-sequencing identifies changes in aged *daf-2* mutants.** (a-b) GO term analysis of whole-worm *daf-2*-regulated genes shows enrichment in stress-resistant genes. (c) Comparison of neuronal and whole-worm Day 8 *daf-2* differentially-expressed genes show high overlap (~30%), but also identify genes specific to neurons and to the whole body. Related to Figure 3h. (d) Comparison of neuronal Day 1 and Day 8 *daf-2* differentially expressed genes identifies a set of consistent and sets of changed genes.

**Figure S6. DAF-16-dependent and -independent *daf-2*-regulated genes show different features.** (a) Volcano plot of *daf-2* vs N2 differentially expressed genes during aging. 1036 genes are more highly expressed in *daf-2* mutants, 1285 genes are higher in N2 ($\log_2$[Fold-change(*daf-2* vs N2)] >0.5, p-adjusted <0.05). (b-c) Neuronal Day 8 *daf-2* vs N2 differentially expressed GO Terms. (d) Tissue query for *daf-2* vs *daf-16;daf-2* differentially expressed genes. (e) Tissue query for *daf-2* vs N2 differentially expressed genes. (f) Full image of comparison of neuronal Day 8 *daf-2* vs *daf-16;daf-2* and *daf-2* vs N2 differentially expressed genes. Related to Figure 5B.

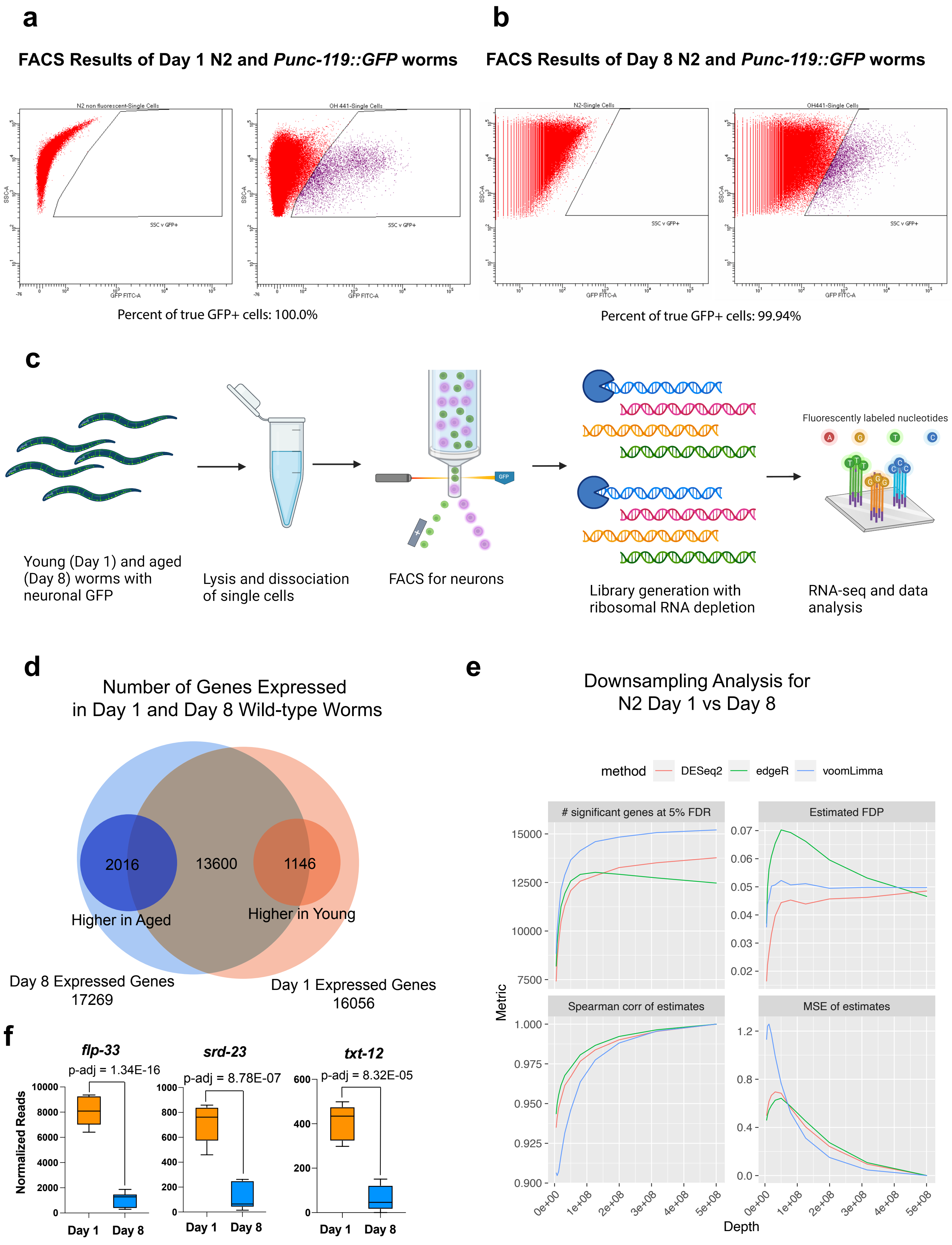


**Supplemental Figure 1**

**Table S1:** Whole-worm DEseq2 results

**Table S2:** Neuronal WT Day 1 vs Day 8 DEseq2

**Table S3:** Neuronal Day 8 *daf-2* vs *daf-16;daf-2* DEseq2

**Table S4:** Neuronal Day 8 *daf-2* vs N2 DEseq2

**Table S5:** Neuronal Day 8 *daf-16;daf-2* vs N2 DEseq2

**Table S6:** Number of sequencing reads

**Table S7:** Raw behavioral data

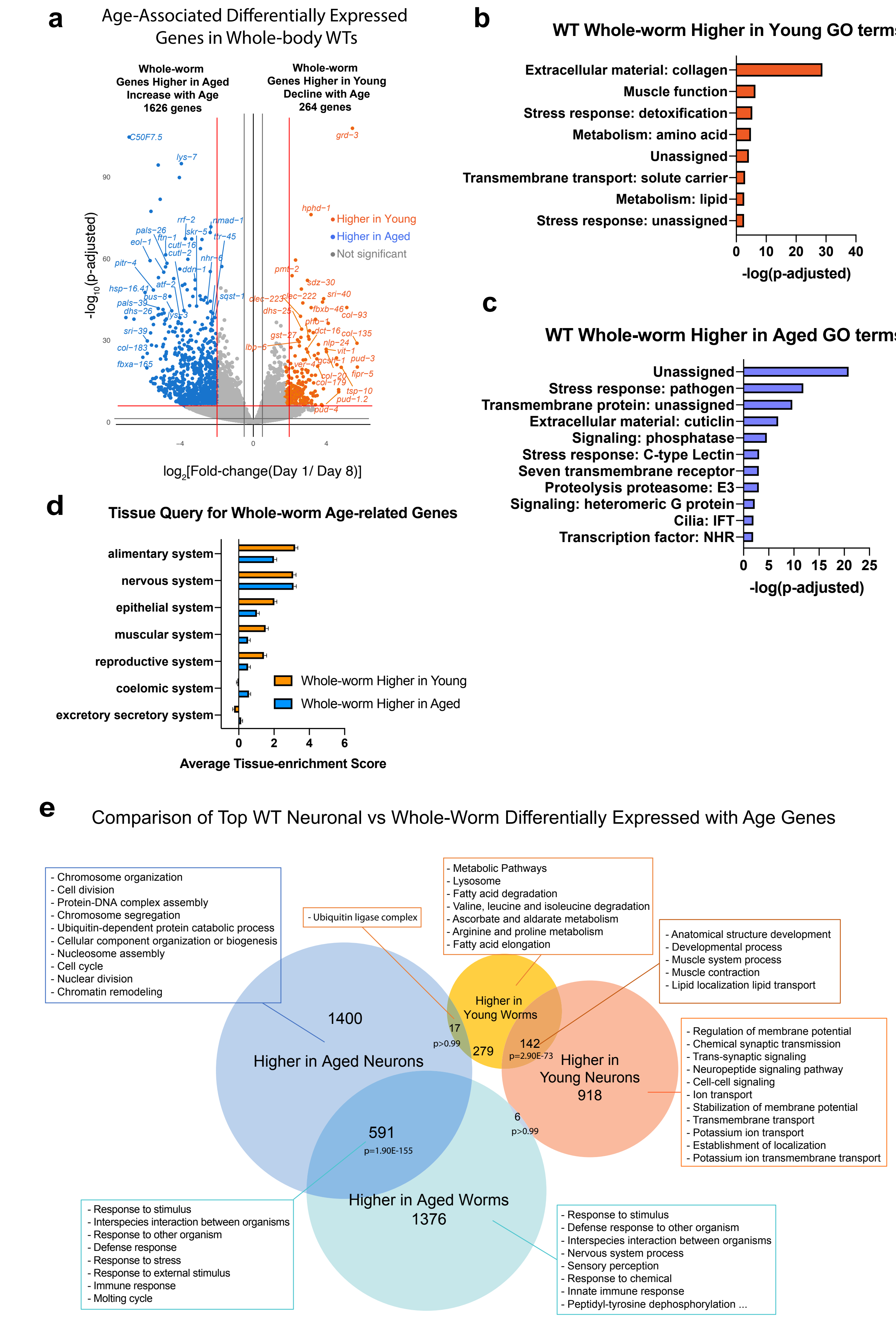


**Supplemental Figure 2**

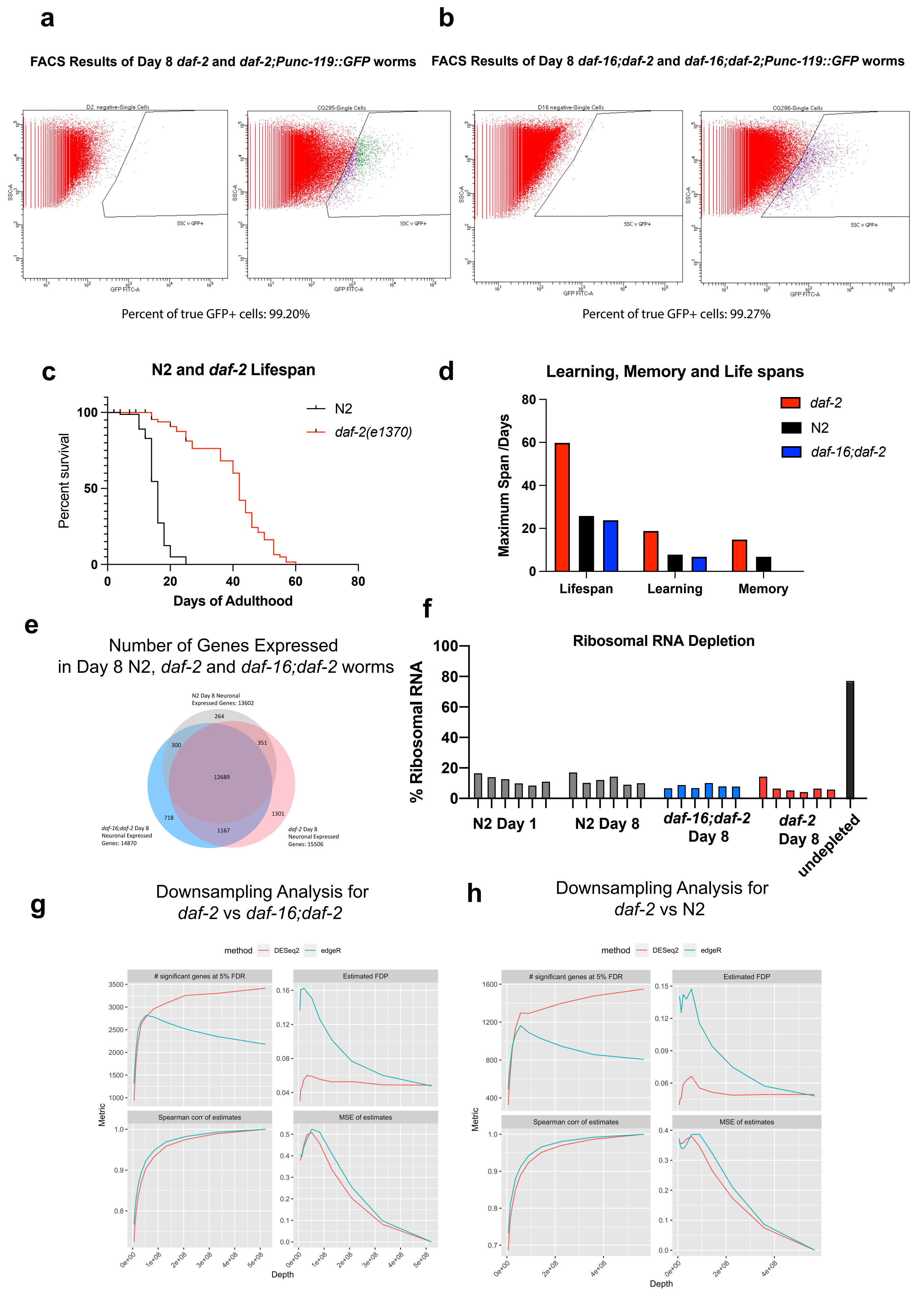


**Supplemental Figure 3**

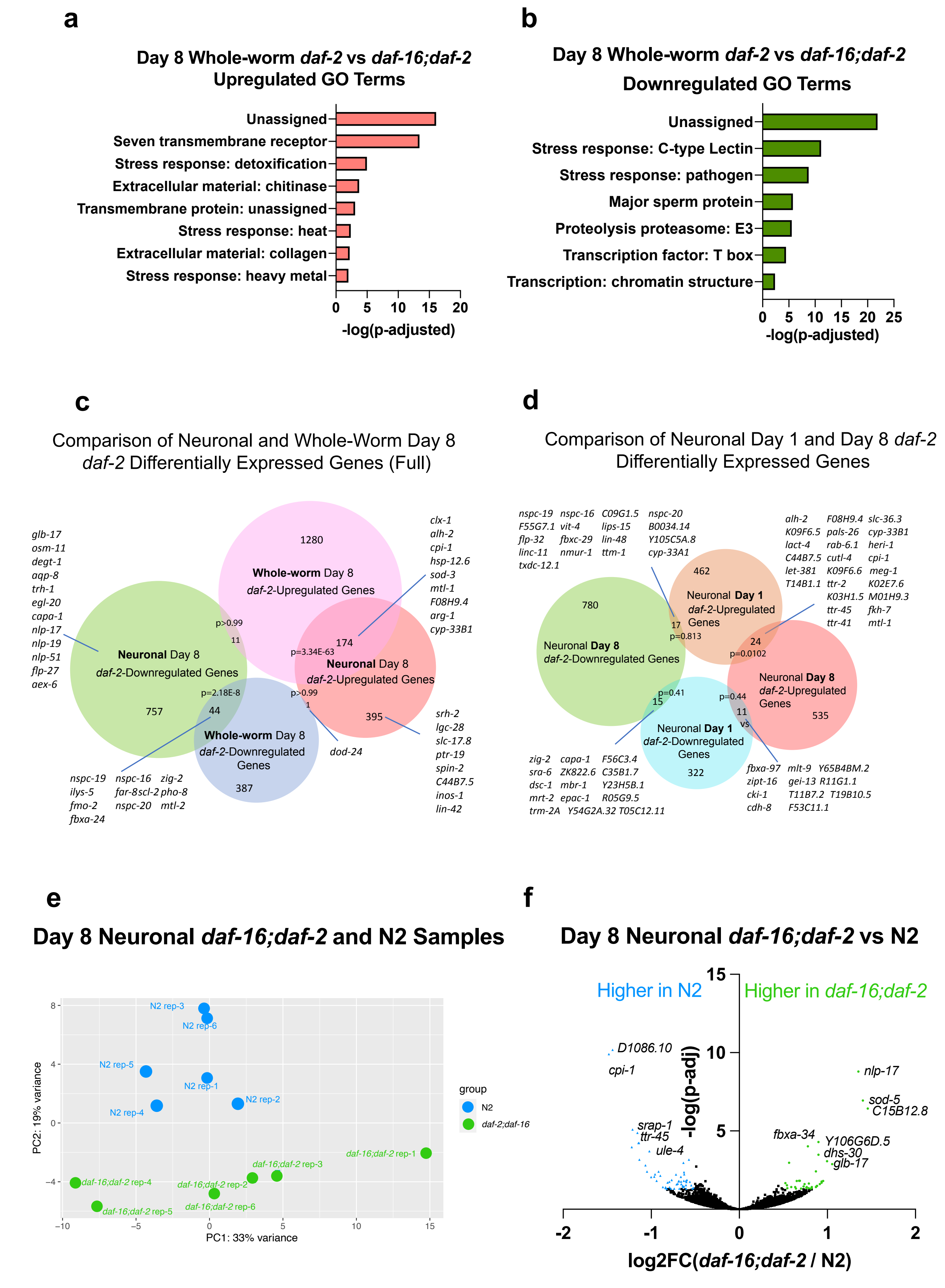


### Supplemental Figure 4

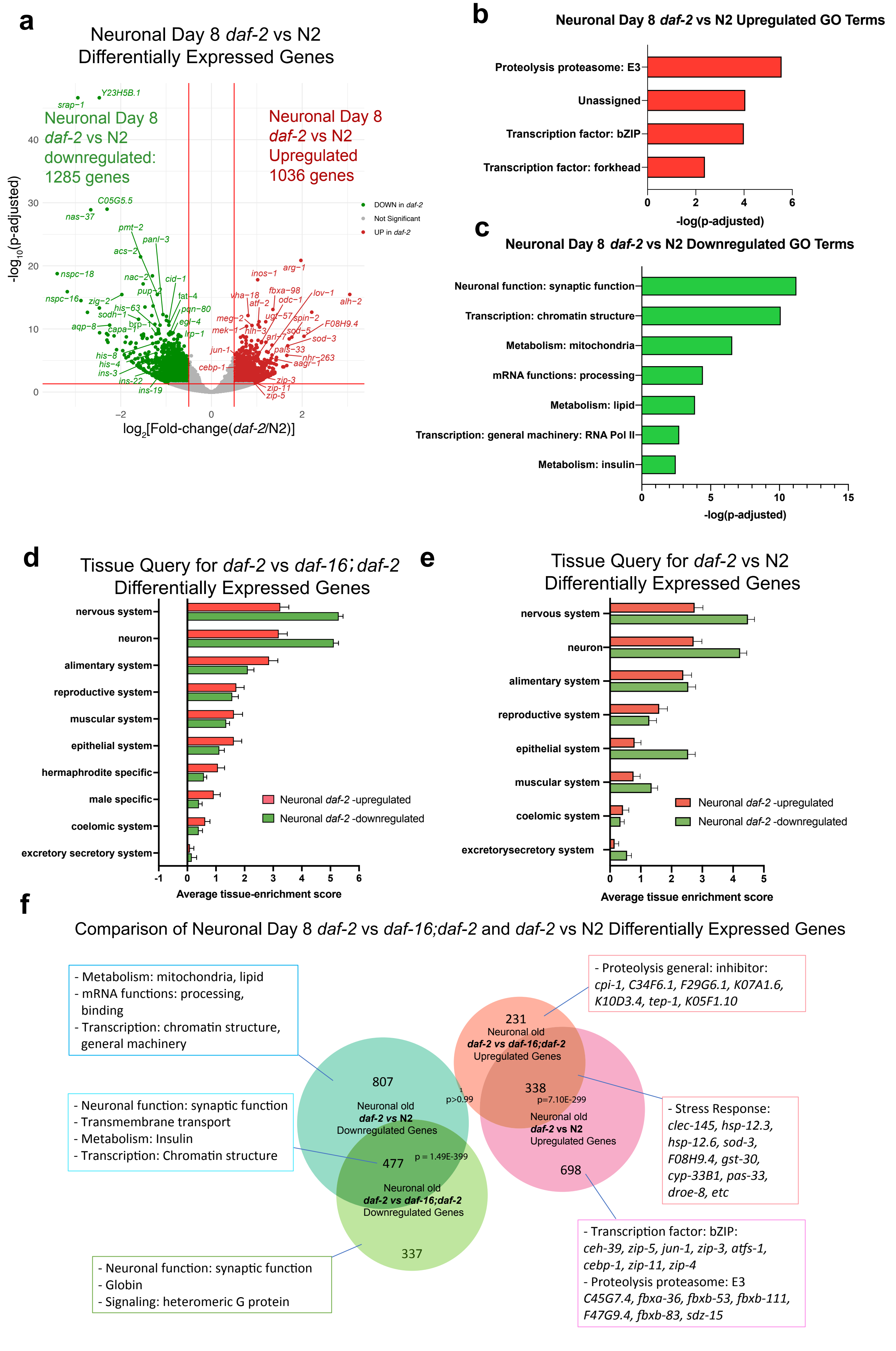


**Supplemental Figure 5**

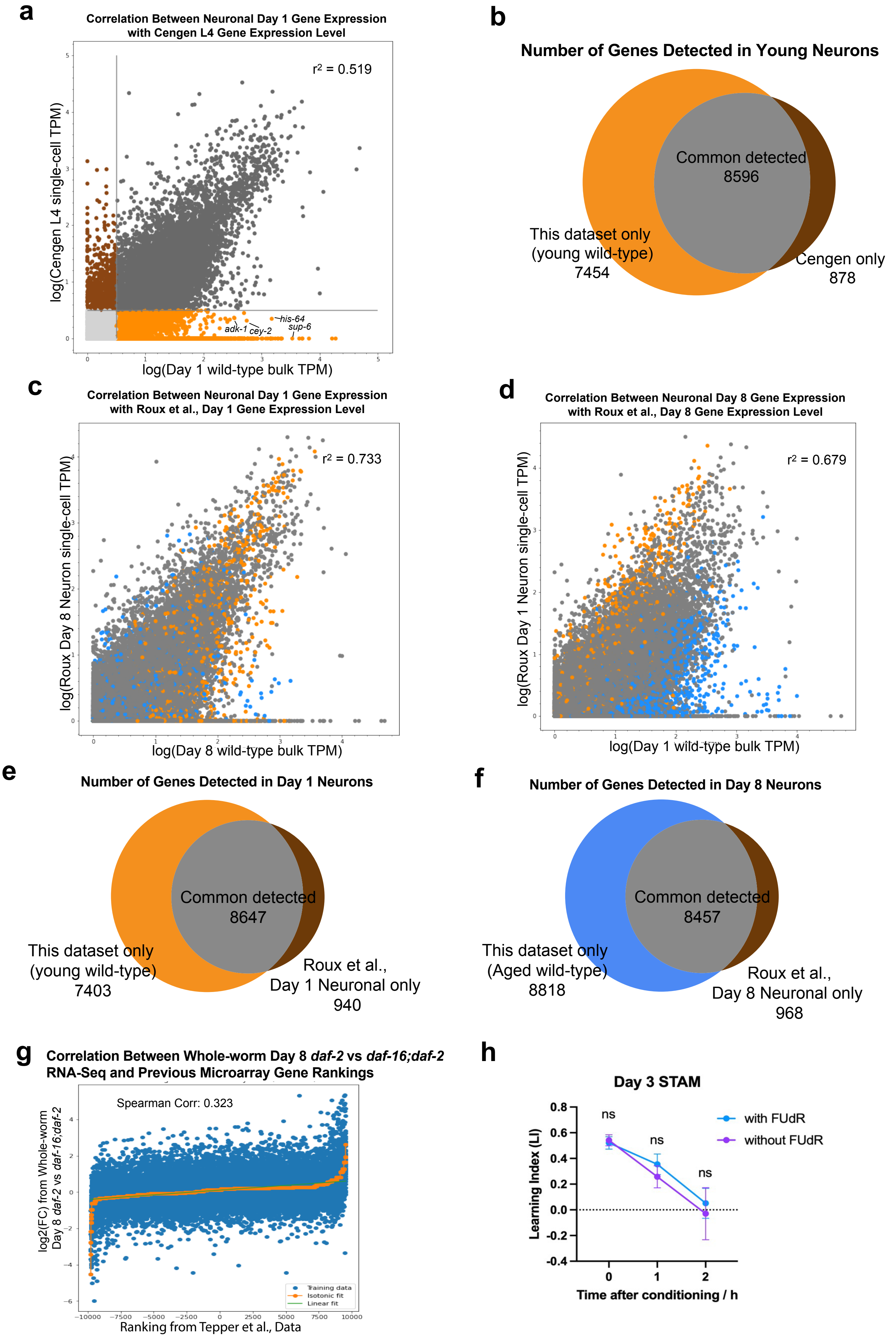


**Supplemental Figure 6**
